# Co-occurrence of VIM-positive non-*aeruginosa Pseudomonas* spp. and *Pseudomonas aeruginosa* in the hospital: A detailed molecular comparison of *bla*_VIM-2_-containing plasmids from environmental and clinical isolates

**DOI:** 10.1101/2024.06.29.601351

**Authors:** Jannette Pirzadian, Willemien H. A. Zandijk, Astrid P. Heikema, Heidy H. H. T. Koene, Wil H. F. Goessens, Margreet C. Vos, Corné H. W. Klaassen, Juliëtte A. Severin

## Abstract

**Background:** *Pseudomonas aeruginosa* is a bacterial pathogen responsible for severe hospital-acquired infections, and is capable of forming persistent reservoirs in hospital sink drains, creating a transmission risk. In our hospital, we have cultured not only carbapenemase-producing, Verona Integron-encoded Metallo-beta-lactamase (VIM)-positive *P. aeruginosa*, but also VIM-positive non-*aeruginosa Pseudomonas* spp., from sink drains. Previously, a *bla*_VIM-2_-containing, conjugative plasmid conferring carbapenem resistance was found in a clinical *P. aeruginosa* isolate in our hospital.

**Objective:** To investigate if other *Pseudomonas* spp. from our hospital also carried a *bla*_VIM-2_-containing plasmid, genetically characterize these plasmids, and compare these plasmids to publicly-available plasmid sequences to identify their source.

**Methods:** Whole-genome sequencing was used to sequence chromosomes and possible plasmids from VIM-positive non-*aeruginosa Pseudomonas* spp. and VIM-positive *P. aeruginosa* strains. Hybrid assemblies were generated to reconstruct plasmid sequences. All isolates were obtained from environmental sampling or clinical cultures during a prolonged VIM-positive *P. aeruginosa* outbreak in our hospital.

**Results:** An identical *bla*_VIM-2_-containing plasmid was found in six non-*aeruginosa Pseudomonas* (*P. carnis*, *P. oleovorans*, and a novel species) and in three *P. aeruginosa* isolates. The previously-reported *bla*_VIM-2_-containing plasmid was found in three additional *P. aeruginosa* isolates. All *P. aeruginosa* belonged to high-risk clones ST111 or ST446. The two plasmids were derived from both clinical isolates and isolates from sink cultures, and were unique to our hospital when compared to other publicly-available plasmid sequences.

**Conclusions:** During a VIM-positive *P. aeruginosa* outbreak in our hospital, two closely-related, *bla*_VIM-2_-containing plasmids co-occurred in multiple non-*aeruginosa Pseudomonas* spp. and in high-risk *P. aeruginosa* clones ST111 and ST446. Non-*aeruginosa Pseudomonas* spp. cultured from either patient or sink samples may be important sources of carbapenem resistance genes, and should not be overlooked during outbreak investigations. Moreover, plasmid analysis is essential to fully understand transmission routes in hospitals.

## 1. Introduction

*Pseudomonas aeruginosa* is a major cause of severe, hospital-acquired infections (De Oliveira *et al*., 2020; Pelegrin *et al*., 2021). It is intrinsically resistant to multiple antimicrobial classes, and the global emergence of carbapenem-resistant *P. aeruginosa* strains, especially belonging to high-risk sequence types (ST) ST111, ST235, and ST175, is of concern (Witney *et al*., 2014; Murray *et al*., 2015; Oliver *et al*., 2015; Cabot *et al*., 2016; Treepong *et al*., 2018). Infections caused by such strains are associated with increased morbidity and mortality to patients in healthcare facilities (Mesaros *et al*., 2007; Oliver *et al*., 2015). Carbapenem resistance in these strains often occurs due to the production of metallo-beta-lactamase (MBL) enzymes (Hong *et al*., 2015). Verona Integron-encoded Metallo-beta-lactamase (VIM) is the most frequently-encountered MBL in *P. aeruginosa* worldwide, and is often encoded in *P. aeruginosa* as a mobile resistance gene cassette localized on a class 1 integron (Hong *et al*., 2015; Oliver *et al*., 2015; Karampatakis *et al*., 2018).

At the Erasmus MC University Medical Centre (Erasmus MC), VIM-positive *P. aeruginosa* clones have emerged since 2003, causing colonisations and infections in patients. Isolates belonged primarily to international, high-risk clones ST446 and ST111 (Van der Bij *et al*., 2011; 2012). Environmental sources and transmission routes have since been investigated, and sinks were identified as a persistent environmental reservoir in our hospital (Voor in ’t holt *et al*., 2018; Pham *et al*., 2022). Although *bla*_VIM_-encoding integrons are usually located on the *P. aeruginosa* chromosome, Van der Zee *et al*. recently reported a *bla*_VIM-2_-containing, conjugative plasmid in *P. aeruginosa* strain S04 90, a ST446 clone isolated from a patient in our hospital in 2009 (Van der Zee et al., 2018). This is worrisome, as plasmids are transferred easily between bacteria of the same species and even within the same genus. During the yearslong outbreak in our hospital, several VIM-positive non-*aeruginosa Pseudomonas* spp. were also recovered from the patient environment and sporadically from patients. In this study, we screened both VIM-positive *P. aeruginosa* and non-*aeruginosa Pseudomonas* isolates for the presence of *bla*_VIM-2_-containing plasmids, genetically characterized plasmids that were found, and compared them to other publicly-available plasmid sequences in order to identify their source.

## 2. Materials and methods

### 2.1 Setting

The Erasmus MC is a large, tertiary-care hospital located in Rotterdam, the Netherlands. Adult intensive care unit 2 (ICU-2) comprised single-occupancy patient rooms or patient isolation rooms with anterooms. The surgical ward comprised two- and four-bed patient rooms, and mainly admitted patients that had undergone digestive tract surgery. The urology ward comprised two-bed, four-bed, and single-occupancy patient rooms.

### 2.2 Selection of isolates

Environmental sampling began in the Erasmus MC in 2011 following recognition of the outbreak. Sink culture samples were obtained using dry swabs (BBL CultureSwab Plus, Copan Italia, Brescia, Italy) on wash basins, countertops, faucet aerators, and sink drains (Pirzadian *et al*., 2023). Swabs were incubated overnight at 35°C in 5 ml of enrichment broth (tryptic soy broth with 2 mg/l ceftazidime and 50 mg/l vancomycin). DNA was extracted from the enrichment broth using the MagNA Pure 96 platform (Roche Diagnostics, Almere, the Netherlands) according to the manufacturer’s instructions. Screening for *bla*_VIM_ was performed using a previously-described real-time PCR protocol (Pirzadian *et al*., 2020). PCR-positive broths were subcultured on blood and MacConkey agars, and MALDI-TOF (Bruker Daltonik, Bremen, Germany) was used for *Pseudomonas spp.* identification. VIM-positive *P. aeruginosa* and non-*aeruginosa Pseudomonas* spp. were stored at -80°C. Additional VIM-positive *P. aeruginosa* and non-*aeruginosa Pseudomonas* isolates were obtained from sink drain samples cultured on diverse media using a microbial culturomics technique (Pirzadian *et al*., 2020). In the current study, all VIM-positive non-*aeruginosa Pseudomonas* isolates were included in the analysis, as well as VIM-positive *P. aeruginosa* that had either already been sequenced in previous studies, or were collected from specific locations in the hospital (e.g. wards and patient rooms considered hotspots for VIM-positive *Pseudomonas* spp.) (Pirzadian *et al*., 2020; 2021).

Clinical cultures were obtained as previously described (Voor in ’t holt *et al*., 2018). The original *P. aeruginosa* S04 90 clinical isolate was included, which was registered at the Erasmus MC under the name VIM-PA R-340. Written approval to use these isolates was received from the medical ethics research committee of the Erasmus MC (MEC-2015-306).

### 2.3 Whole-genome sequencing and plasmid reconstruction

DNA was isolated from freshly-grown colonies using the Zymo Quick-DNA Fungal/Bacterial Miniprep Kit (Zymo Research, Irvine, CA, USA). Libraries were prepared using the Nextera DNA Flex Library Prep as recommended by the manufacturer (Illumina, San Diego, CA, USA), then sequenced using Illumina technology to generate 2×150 bp paired-end reads and >30x coverage. *De novo* assemblies were created using CLC Genomics Workbench v21 (Qiagen, Hilden, Germany) with default parameters. The identity of non-*aeruginosa Pseudomonas* strains was established by analysing genomic assemblies using the online Type Strain Genome Server (TYGS) platform (https://tygs.dsmz.de/) (Meier-Kolthoff and Göker, 2019). *P. aeruginosa* multi-locus sequence types (MLST) were determined using the MLST plugin in BioNumerics v7.6 software (Applied Maths, St-Martens-Latem, Belgium) synchronized to the PubMLST.org database (https://pubmlst.org/).

To identify strains carrying identical plasmids, a resequencing approach was used by mapping the Illumina reads of the sample to the respective plasmid sequences in CLC Genomics Workbench. Standard read mapping parameters were adjusted from 50% identity and 80% coverage to a highly stringent 98%. The resulting mapping graphs were checked for structural integrity and/or loss of coverage. For selected strains, plasmid identification and reconstruction were established by also generating hybrid assemblies.

*De novo* reconstruction of plasmid sequences was performed by first isolating high molecular weight DNA with the Qiagen Genomic-tip 20/G protocol (Qiagen). Libraries were constructed using the Rapid Barcoding kit (Oxford Nanopore Technologies (ONT), Oxford, United Kingdom), and sequenced on a MinION R-9.4 flowcell (ONT). The combination of MinION and Illumina sequences were used to create hybrid assemblies in Unicycler v0.5.0 (Wick *et al*., 2017). For easier comparison, the starting position and orientation from the reconstructed circular plasmid sequences were adjusted to match that of the previously-described plasmid sequence from *P. aeruginosa* strain S04 90 (Van der Zee *et al*., 2018).

Whole-genome alignments were made in CLC Genomics Workbench. A global comparison was made using a minimum initial seed length (ISL) of 25 and a minimum alignment block length (ABL) of 100. For a more high-resolution comparison, parameters were adjusted to an ISL of 5 and an ABL of 25. Syntenic regions were indicated with colours and their identity was determined by pairwise BLAST analysis (https://blast.ncbi.nlm.nih.gov/Blast.cgi). A k-mer-based unrooted neighbour joining tree (k=9) was constructed using the Mahalanobis distance measure in CLC Genomics Workbench. Identification of insertion sequences (IS) was performed using the web-based ISfinder software (https://isfinder.biotoul.fr/blast.php) with default parameters (Siguier *et al*., 2006).

## 3. Results

### 3.1 Reconstruction of the *P. aeruginosa* S04 90 plasmid sequence

Firstly, the original *P. aeruginosa* strain S04 90, stored at the Erasmus MC as VIM-PA R-340, was re-sequenced and the plasmid reconstructed using a hybrid assembly. Our results confirmed the presence and identity of the plasmid sequence with some minor differences: (i) A total of 7 (out of 159,166) nucleotides differed when compared to the published sequence (99.996% identity), and (ii) while the last 21 nucleotides of the published sequence are an exact copy of the first 21 nucleotides, during our reconstruction, only one copy of these sequences was found (Van der Zee *et al*., 2018).

### 3.2 Non-*aeruginosa Pseudomonas* isolates and plasmid sequence analysis

Six isolates of VIM-positive non-*aeruginosa Pseudomonas* spp. contained an identical plasmid sequence: One *P. oleovorans*, two *P. carnis*, and three isolates of a novel *Pseudomonas* sp (Table 1). The two *P. carnis* isolates had been originally identified as *P. alcaligenes* and *P. gessardii*, respectively, by MALDI-TOF, but were later reclassified as *P. carnis* after sequencing. The three isolates of a novel *Pseudomonas* sp. had been originally identified as *P. putida* by MALDI-TOF. The plasmid sequence of all six non-*aeruginosa Pseudomonas* strains also contained the *bla*_VIM-2_ gene. Five strains had been cultured from the sink environment (either wash basin or drain) in the same ward, ICU-2, but from different patient rooms between 2013 and 2016; the sixth strain (a novel *Pseudomona*s sp.) had been isolated from a patient staying in ICU-2 in 2014. No clinical or environmental VIM-positive non-*aeruginosa Pseudomonas* spp. had been isolated from other wards.

**Table 1.** *Pseudomonas* isolates analysed in this study. *P. carnis* strain VIM-PC R-7810 was the earliest non-*aeruginosa Pseudomonas* isolate found carrying the *bla*_VIM-2_-encoding plasmid in our hospital, and as this plasmid sequence differed from the previously-reported *P. aeruginosa* plasmid sequence, in the column “Plasmid identity,” isolates are indicated with a shaded box as having an identical plasmid sequence to either *P. aeruginosa* strain VIM-PA R-340 or to *P. carnis* strain VIM-PC R-7810. n.a., Not available.

| Isolate name | Species | Sequence type | Year of isolation | Isolation source | Ward | Plasmid identity |  |
| --- | --- | --- | --- | --- | --- | --- | --- |
|  |  |  |  |  |  | VIM-PA R-340 | VIM-PC R-7810 |
| VIM-PA R-340 | *P. aeruginosa* | 446 | 2009 | Clinical | Urology |  |  |
| VIM-PC R-7810 | *P. carnis* | n.a. | 2013 | Environmental | ICU-2 |  |  |
| VIM-PO R-9888 | *P. oleovorans* | n.a. | 2014 | Environmental | ICU-2 |  |  |
| VIM-PSp R-9374 | Novel<br>*Pseudomonas*<br>*sp.* | n.a. | 2014 | Environmental | ICU-2 |  |  |
| VIM-PSp R-9773 | Novel<br>*Pseudomonas*<br>*sp.* | n.a. | 2014 | Clinical | ICU-2 |  |  |
| VIM-PSp R-12053 | Novel<br>*Pseudomonas*<br>*sp.* | n.a. | 2016 | Environmental | ICU-2 |  |  |
| VIM-PC 49337 | *P. carnis* | n.a. | 2016 | Environmental | ICU-2 |  |  |
| VIM-PA 43337 | *P. aeruginosa* | 111 | 2016 | Environmental | ICU-2 |  |  |
| VIM-PA 5009 | *P. aeruginosa* | 111 | 2017 | Environmental | ICU-2 |  |  |
| VIM-PA R-14745 | *P. aeruginosa* | 111 | 2017 | Environmental | ICU-2 |  |  |
| VIM-PA R-15694 | *P. aeruginosa* | 111 | 2017 | Clinical | ICU-2 |  |  |
| VIM-PA R-16609 | *P. aeruginosa* | 111 | 2018 | Environmental | Surgery |  |  |
| VIM-PA R-16738 | *P. aeruginosa* | 111 | 2018 | Environmental | Surgery |  |  |

An alignment of the plasmid sequences from *P. aeruginosa* strain VIM-PA R-340 and *P. carnis* strain VIM-PC R-7810 was performed, and revealed a high general level of similarity with minor differences between the two sequences (Figure 1). The most obvious difference was the rearrangement of a 4.7 kbp region encoding a hypothetical protein (Figure 1, box A). Additionally, a 2.6 kbp insertion sequence 99% identical to ISpst3 (IS21 family) was identified in the plasmid sequence of VIM-PA R-340, and only in the chromosome of VIM-PC R-7810 (Figure 1, box B). This insertion sequence encoded an integrase, ATPase AAA, and contained a 5 bp inverted repeat (IR) sequence but no characteristic direct repeat (DR) sequence. In the corresponding region of the VIM-PC R-7810 plasmid, a 3.3 kbp region with a replicative DNA helicase and glyoxylase-encoding gene was present instead. Finally, a 1.2 kbp insertion sequence 98% identical to IS222 (IS3 family), encoding two transposes and containing a 3 bp DR sequence and 9 bp IR element, was only found in the plasmid sequence of VIM-PA R-340 (Figure 1, box C). Apart from these differences, the two plasmid sequences were identical, including a class 1 integron on which the *bla*_VIM-2_ gene was encoded. Moreover, the sequence of the integron was identical to the recently-described *bla*_VIM-2_-encoding class 1 integron carried by the majority of VIM-2-positive *P. aeruginosa* isolates (i.e. Group 1) in the Netherlands (Pirzadian *et al*., 2021).

**Figure 1.**
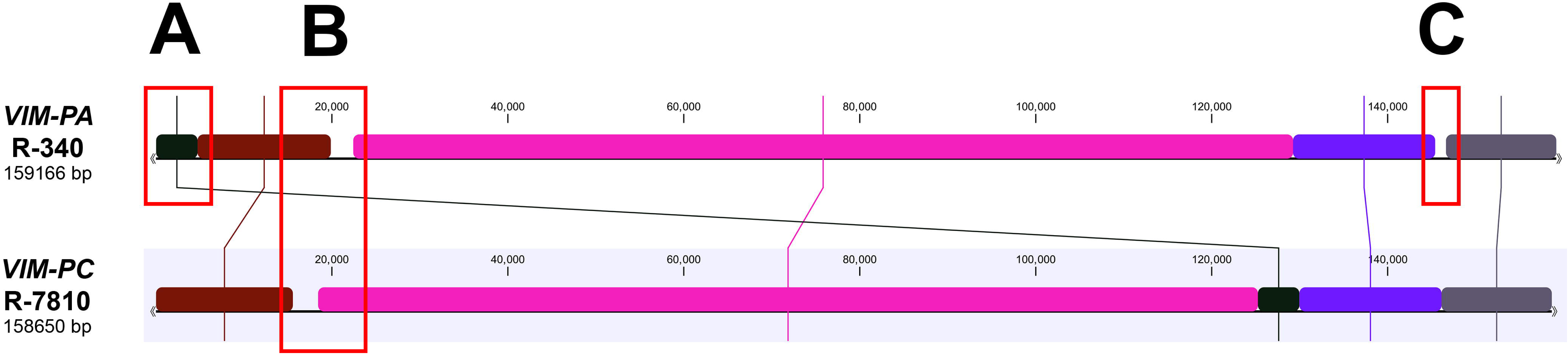
Alignment between the plasmid sequences from *P. aeruginosa* strain VIM-PA R-340 and *P. carnis* strain VIM-PC R-7810. All non-*aeruginosa Pseudomonas* spp. in this study shared 100% sequence identity to the *P. carnis* plasmid. Boxes A, B, and C indicate differences between the two plasmid sequences, which are detailed further in the text.

To place these plasmids into an international context, a BLAST search was performed, and fifteen plasmid sequences with >50% coverage to the plasmid sequence from VIM-PA R-340 were retrieved and subjected to a whole-genome alignment. All included plasmids shared a highly similar backbone structure (Figure 2). However, k-mer-based clustering showed that the plasmids were still quite diverse; the two plasmids identified in our study clustered closely together, and were distinct from the other NCBI-deposited plasmid sequences (Figure 3).

**Figure 2.**
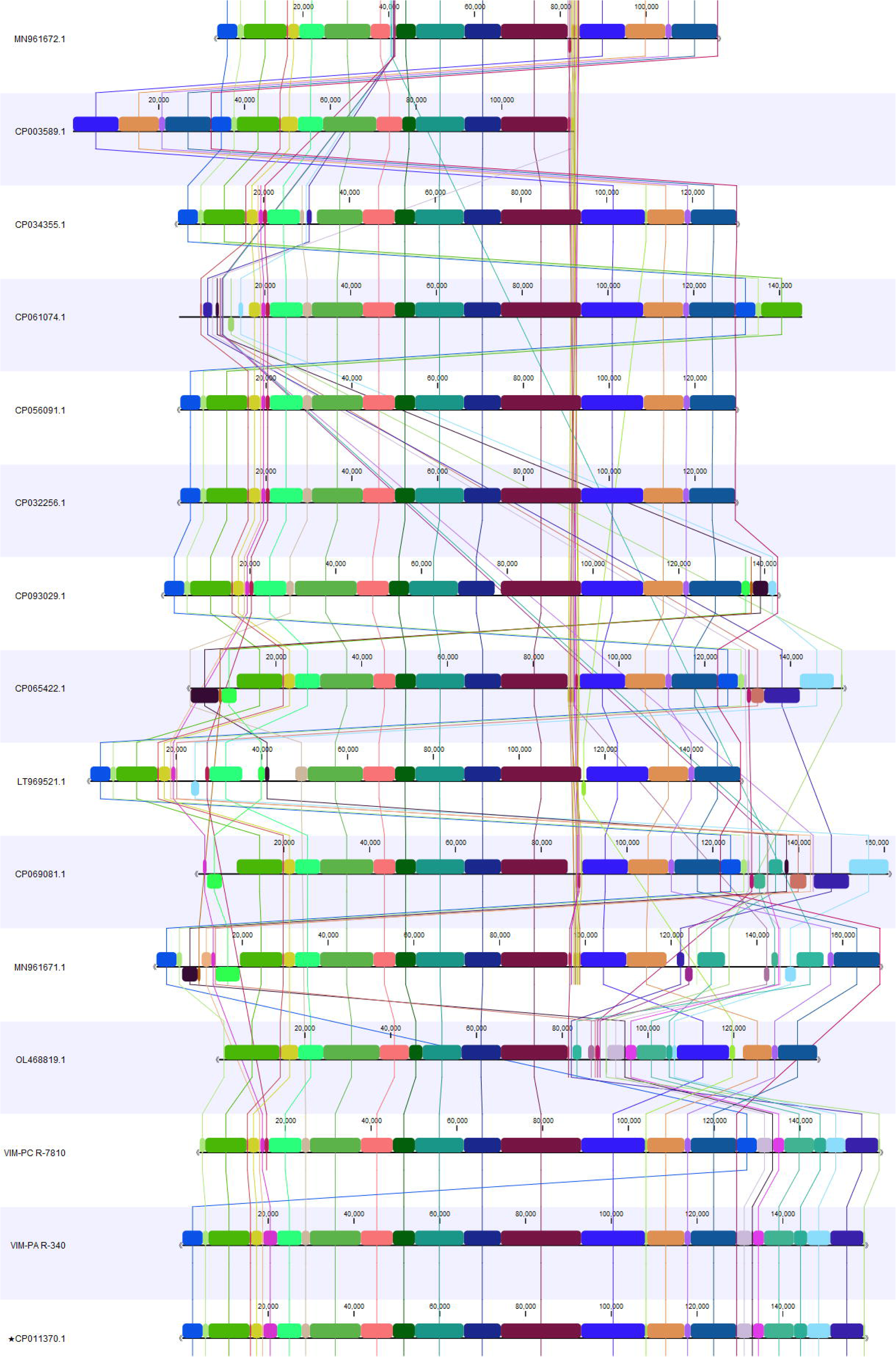
Whole-genome alignment between fifteen plasmid sequences identified using NCBI’s BLAST showing a similar backbone structure. The sequences of the published *P. aeruginosa* strain S04 90 (accession no. CP011370.1, indicated with an asterisk) (Van der Zee *et al*., 2018), *P. aeruginosa* strain VIM-PA R-340, and *P. carnis* strain VIM-PC R-7810 are included in the figure. Double arrows (<< >>) are used to indicate closed circular constructs.

**Figure 3.**
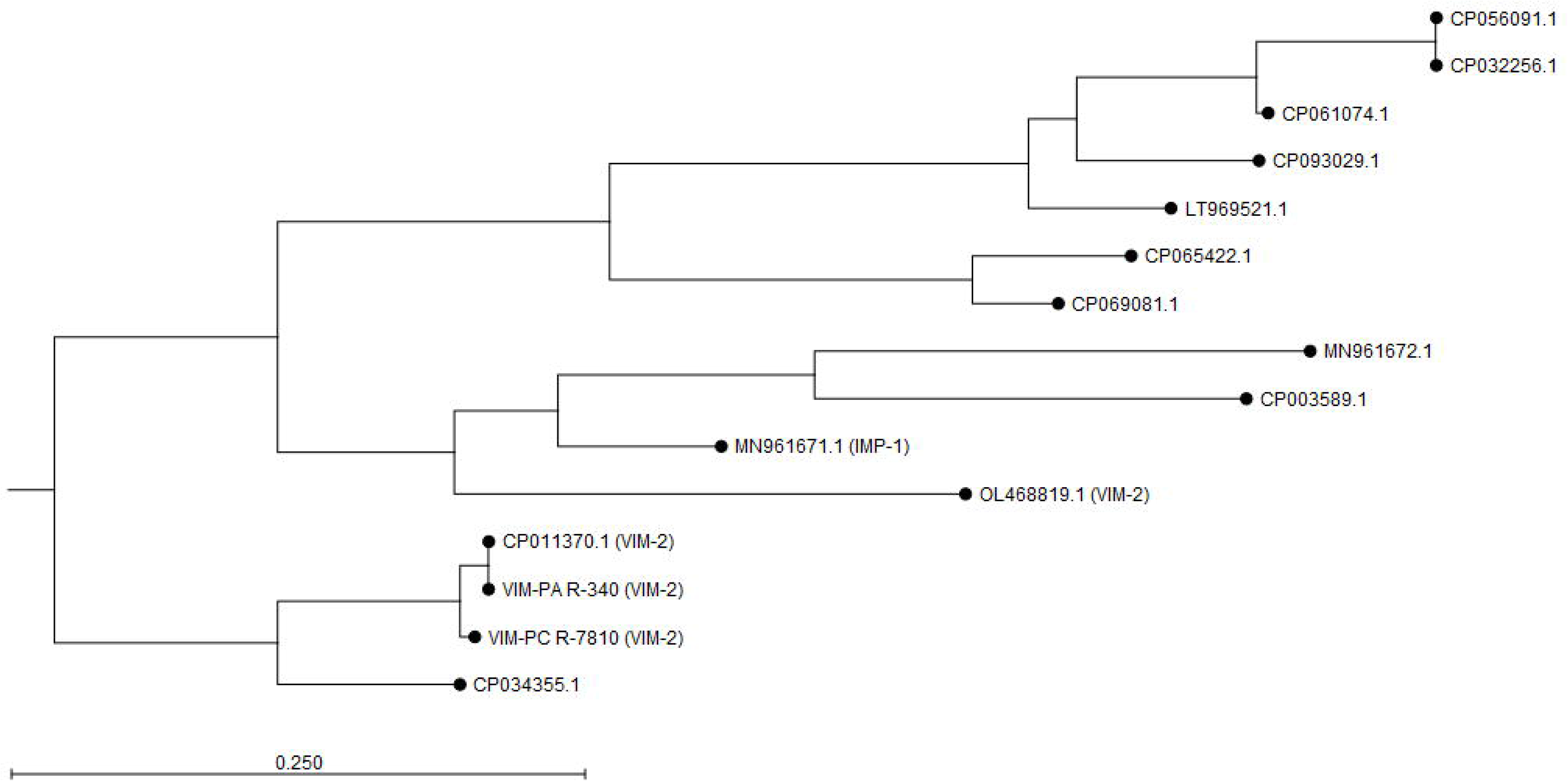
Unrooted neighbour joining tree of the fifteen plasmid sequences identified using NCBI’s BLAST. The published *P. aeruginosa* strain S04 90 (accession no. CP011370.1) (Van der Zee *et al*., 2018), *P. aeruginosa* strain VIM-PA R-340, and *P. carnis* strain VIM-PC R-7810 are shown closely clustered at the bottom of the tree. If a plasmid sequence contained a carbapenemase gene, this is indicated after the accession number or strain name in parentheses. IMP, Imipenemase; VIM, Verona Integron-encoded Metallo-beta-lactamase.

### 3.3 *P. aeruginosa* isolates and plasmid sequence analysis

Three VIM-positive *P. aeruginosa* isolates contained an identical plasmid sequence to the plasmid from *P. aeruginosa* strain VIM-PA R-340: VIM-PA 43337, VIM-PA 5009, and VIM-PA 14745 (Table 1). These *P. aeruginosa* strains were isolated between 2016 and 2017 from the same patient room in ICU-2, and were all obtained from sink drain samples. *P. aeruginosa* strain S04 90 / VIM-PA R-340 had been isolated in 2009 from a patient admitted to ICU-2, but from a different patient room.

Three VIM-positive *P. aeruginosa* strains contained an identical plasmid sequence to the plasmid from *P. carnis* strain VIM-PC R-7810: VIM-PA R-15694, VIM-PA R-16609, and VIM-PA R-16738. *P. aeruginosa* VIM-PA R-15694 was isolated from a patient in 2017 admitted to the ICU-2 patient room where *P. carnis* VIM-PC R-7810 had been isolated in 2013. On the sampling date of the clinical culture, the patient had been transferred to the surgical ward; in 2018, the two other *P. aeruginosa* strains were isolated from sink drain samples from the same surgical ward patient room.

All six VIM-positive *P. aeruginosa* strains were identified as ST111 high-risk clones. *P. aeruginosa* strain VIM-PA R-340 was confirmed to be high-risk clone ST446. Additionally, the *bla*_VIM-2_ gene was only found in plasmid sequences and not in the chromosome of any of these strains.

## 4. Discussion

This study reinforces findings from a growing body of literature that non-*aeruginosa Pseudomonas* spp. may act as reservoirs for the *bla*_VIM-2_ gene, and that the hospital environment may be involved in the circulation of such genes. We identified a *bla*_VIM-2_-containing plasmid at the Erasmus MC carried by six non-*aeruginosa Pseudomonas* isolates (*P. oleovorans*, *P. carnis*, and a novel *Pseudomonas* sp.) and three *P. aeruginosa* ST111 isolates. This plasmid was highly similar to a second, previously-described plasmid from *P. aeruginosa* strain S04 90, a ST446 clone, found in our hospital (Van der Zee *et al*., 2018). We identified three other *P. aeruginosa* ST111 isolates with plasmids identical to this previously-described plasmid. Isolates were obtained from both clinical and sink cultures. The two plasmids found in this study were genetically distinct from other publicly-deposited plasmid sequences, so the source of the plasmids remain unknown. However, it is remarkable that the two plasmids were closely related, and remained stable for the entire study period spanning nine years. We hypothesize that these two *bla*_VIM-2_-encoding plasmids likely transferred between high-risk *P. aeruginosa* clones and other *Pseudomonas* spp., using sink drains as a primary environmental reservoir, within the Erasmus MC. To our knowledge, this is also the first report of VIM-2-positive *P. carnis*, and the second report of *P. oleovorans* containing a *bla*_VIM-2_-encoding plasmid.

*P. aeruginosa* is frequently recovered from wet niches in the hospital environment, such as in sink drains and showers, where it may form a persistent reservoir capable of dispersing viable cells within droplets to the patient’s innate environment (Hota *et al*., 2009; Breathnach *et al*., 2012; Hopman *et al*., 2019). Other *Pseudomonas* spp., such as members of the *P. putida* group, may also be present in these niches; in fact, we confirmed in a prior study that in our hospital, VIM-positive non-*aeruginosa Pseudomonas* spp. formed part of the microbial consortium in sink drain biofilms containing high-risk VIM-positive *P. aeruginosa* clone ST111 (Pirzadian *et al*., 2020). Nonetheless, non-*aeruginosa Pseudomonas* spp. are rarely the sources of human infections, so are often not studied or surveilled in a clinical setting. However, numerous reports have identified non-*aeruginosa Pseudomonas* spp. as carriers of MBL genes, which our results also support (Quinteira, Ferreira, and Peixe, 2005; Juan *et al*., 2010; Scotta *et al*., 2011; Peter *et al*., 2017; Büchler *et al*., 2022). Unique to our study was the strong involvement of sink drains in the long-term persistence of isolates containing the *bla*_VIM-2_-encoding plasmids, which could be linked to both patient-to-environment and environment-to-patient spread. This also reinforces a previous surveillance study which found that wastewater networks were potential reservoirs for plasmids capable of spreading resistance genes to patient populations (Weingarten *et al*., 2018). To date, MBL-carrying plasmids have been identified more frequently in non-*aeruginosa Pseudomonas* spp. than in *P. aeruginosa*. Our findings underscore that non-*aeruginosa Pseudomonas* spp. should not be ignored during outbreak investigations, but also be included and screened for carbapenemases.

*P. carnis* is a newly-described environmental species involved in the food spoilage of meat and soy products, but it has never before been reported to cause human infection or carry *bla*_VIM_ (Lick *et al*., 2020; De León *et al*., 2021). *P. oleovorans* is considered an environmental isolate, and in only rare cases has caused infections in immunocompromised individuals (Faccone *et al*., 2014; Gautam *et al*., 2015). In Argentina, researchers discovered a plasmid carried by a *P. oleovorans* clinical isolate containing VIM-2 on a class 1 integron; this integron was recovered from plasmids of different sizes from other non-*aeruginosa Pseudomonas* spp. isolated in different Argentinian hospitals (Faccone *et al*., 2014). This differs from our study in that we found only one plasmid in different non-*aeruginosa Pseudomonas* spp., but their results showed that transmission of a *bla*_VIM-2_-encoding integron among different plasmids was also possible. *P. putida* has also rarely been a source of opportunistic human infections, but remains the most frequently-reported non-*aeruginosa Pseudomonas* to carry multidrug-resistance genes, including *bla*_VIM_ (Juan *et al*., 2010; Marchiaro *et al*., 2014; Peter *et al*., 2017). However, our study also showed that isolates identified as *P. putida* may not always be true *P. putida*, but may belong to other species, including novel species; limitations to MALDI-TOF meant that sequencing was required for this distinction. Nevertheless, our findings emphasize that multiple non-*aeruginosa Pseudomonas* spp. are capable of acting not only as MBL reservoirs, but also as reservoirs for plasmids and integrons.

Using a BLAST search and aligning matching plasmid sequences, we identified similar backbone structures in thirteen NCBI-deposited sequences when compared to *P. aeruginosa* VIM-PA R-340 and *P. carnis* VIM-PC R-7810. However, a k-mer-based analysis highlighted that the plasmids in our study were still genetically distinct from these other sequences, so the source of our two plasmids could not be identified. The nearest neighbour (accession no. CP034355.1) was a plasmid sequence from a *P. aeruginosa* strain that carried the carbapenemase gene *bla*_IMP-13_ on its chromosome; neither *bla*_VIM-2_ nor another carbapenemase were encoded on this plasmid. Two more distantly-related strains also carried carbapenemase genes, but all of the other public sequences lacked a carbapenemase gene. These differences suggest that the plasmids in our study are unique to our hospital, and in wider context, are two *Pseudomonas* plasmids unique to the Netherlands. Additionally, while *bla*_VIM_ is typically encoded on the *P. aeruginosa* chromosome, for all isolates in this study, *bla*_VIM_ was only located on the plasmid. Therefore, we hypothesize that at some point, a class 1 integron was mobilized onto a plasmid, which then diversified among *Pseudomonas* strains within the Erasmus MC, most likely within the wet biofilms in sink drains.

A strength of this study was that all VIM-positive non-*aeruginosa Pseudomonas* isolates were sequenced to screen for the *bla*_VIM-2_-encoding plasmid. Additionally, while it was not routinely performed, extensive environmental sampling took place at the Erasmus MC since 2011, and strains were kept in long-term storage; this was beneficial in tracking the movement and evolution of the *bla*_VIM-2_-encoding plasmids over the years. However, this study also had some limitations. Firstly, not all VIM-positive *P. aeruginosa* strains were sequenced. Strains were selected based on their year of isolation and their location of origin (e.g. wards/patient rooms that were hotspots for VIM-positive *P. aeruginosa* during the outbreak, or where VIM-positive non-*aeruginosa Pseudomonas* spp. were isolated), or if their sequences were already available. Secondly, environmental sampling was not routinely performed in the hospital; the decision to swab the environment was guided by patients becoming unexpectedly positive for a multidrug-resistant microorganism(s)/suspected transmission, or to monitor the effects of interventions designed to limit dispersal from VIM-positive *P. aeruginosa* sink drain reservoirs (Pirzadian *et al*., 2023). Therefore, other *Pseudomonas* strains carrying the *bla*_VIM-2_-containing plasmid may have been missed. Finally, conjugation experiments were not performed using the two identified plasmids to verify transmissibility. This was performed in the article by Van der Zee *et al*., which showed that the plasmid of *P. aeruginosa* strain S04 90 was conjugative (Van der Zee *et al*., 2018); our plasmids were highly similar to this published plasmid, so it is likely they are also conjugative. Moreover, the presence of identical sequences in multiple *Pseudomonas* spp. strongly suggests plasmid transfer.

## 5. Conclusions

Our results revealed the co-occurrence of two closely-related *bla*_VIM-2_-containing plasmids in multiple non-*aeruginosa Pseudomonas* spp. and in high-risk *P. aeruginosa* clones derived from patient room sinks but also from clinical cultures. This provides evidence that environment-to-patient or patient-to-environment transmission had most likely occurred within our hospital, and that non-*aeruginosa Pseudomonas* spp. played a role in the circulation of the plasmid, though the origin of the plasmid is still unknown. For effective infection prevention and control, non-*aeruginosa Pseudomonas* spp. should be taken into account as potential reservoirs for *bla*_VIM_ or other carbapenemase genes. Moreover, to fully understand transmission routes of resistance genes in the hospital, plasmid analysis is essential.

## Acknowledgements

The authors would like to thank the employees of the Unit Infection Prevention and the Unit Diagnostics from the Erasmus MC department of Medical Microbiology and Infectious Diseases for collecting and storing isolates. The authors would also like to thank Elyne Clyncke for collecting and storing isolates from pilot culturomics experiments. Findings from this study were presented at ASM Microbe 2019 on 23 June 2019 in San Francisco, CA, USA (Poster P605), and published in the abstract book of the 30^th^ edition of ECCMID (Abstract 4559).

## CRediT authorship contribution statement

CK and JS designed and initiated the study. JP and HK performed environmental sampling and gathered isolates. WZ and AH performed all sequencing experiments. CK and WZ assembled and analysed the sequencing data. JP, AH, WG, MV, CK, and JS curated and analysed all other data. JP wrote the manuscript. All authors read and approved the final manuscript.

## Data availability

Sequence data on the isolates in this paper have been deposited in the NCBI SRA database under BioProject PRJNA975904 and BioSample accession numbers SAMN35347199 through SAMN35347207.

